# Actin assembly-inducing protein ActA promotes FcγRIa-mediated *Listeria* internalization

**DOI:** 10.1101/175117

**Authors:** Sofya S. Perelman, Michael E. Abrams, Neal M. Alto

## Abstract

*Listeria monocytogenes* is a Gram-positive intracellular pathogen and the causative agent of human listeriosis. While the ability of *L. monocytogenes* to enter and survive in professional phagocytes is critical to establish a successful infection, mechanisms of invasion are poorly understood. Our previous investigation into the role of type I interferon-stimulated genes in bacterial infection revealed that the human immunoglobulin receptor FcγRIa served as a *L. monocytogenes* invasion factor. FcγRIa-mediated *L. monocytogenes* entry occurred independently of immunoglobulin interaction or bacterial internalins. However, the bacterial determinants that mediate FcγRIa interaction remain unclear. Using a comparative genomics approach, we identify actin assembly-inducing protein ActA as a pathogen specific ligand of FcγRIa. FcγRIa enhanced entry of pathogenic *L. monocytogenes* and *L. ivanovii* strain but not non-pathogenic *L. innocua*. We found that the major virulence regulator PrfA is required for pathogen entry into FcγRIa-expressing cells and identify its gene target *actA* as the critical *Listeria* ligand. ActA alone was sufficient to promote entry into FcγRIa-expressing cells, and this function is independent of its actin nucleating activity. Together, these studies present an unexpected role of ActA beyond its canonical function in actin-based motility and expand our understanding of *Listeria* strategies for host cell invasion.

**Importance:** *Listeria monocytogenes* is a food-borne bacterial pathogen and a causative agent of listeriosis with up to 50% mortality rate in immunocompromised individuals. While the mechanisms of *Listeria* entry into non-phagocytic cells have been extensively characterized, the details of phagocytic cell invasion are still poorly understood. We have recently discovered that human immunoglobulin receptor FcγRIa mediates *Listeria* uptake by monocytic cells. This process occurred independently of canonical immunoglobulin interactions as well as classic *Listeria* internalization factors. Importantly, molecular determinants of *Listeria*-FcγRIa interaction leading to bacterial entry, remained unknown. In this study, we demonstrate that *Listeria* virulence factor actin-assembly inducing protein ActA is required for FcγRIa-mediated entry. Further, ActA was found to be sufficient for the internalization, suggesting its role as a bacterial ligand of FcγRIa. Together, these findings expand our knowledge of mechanisms that *Listeria* has evolved to exploit cellular signaling pathways and immune defense of the host.

## Introduction

Intracellular bacterial pathogens have evolved a variety of strategies to facilitate entry into host cells, which provide a rich source of nutrients and protect from extracellular host defense and clearance (1). Bacterial uptake by non-professional phagocytic cells occurs primarily through zipper and trigger-based mechanisms. “Zippering” takes place when a bacterial surface protein interacts with a specific host cell receptor, resulting in cytoskeletal rearrangements, vacuole formation, and subsequent engulfment of the pathogen. Bacteria employing a trigger mechanism bypass the interaction with host surface receptors. Instead, bacterial secretion systems deliver effector proteins across cellular membranes, directly engaging intracellular machinery of the host to regulate actin dynamics and promote invasion by macropinocytosis. While these mechanisms encompass general strategies of invasion by pathogens, each bacterial species encodes unique repertoires of virulence factors to co-opt host internalization pathways.

The opportunistic intracellular pathogen *Listeria monocytogenes* (*Lm*) is one of the most well-characterized bacteria that uses the zipper mechanism to enter non-phagocytic epithelial cells and hepatocytes (2). The major *Lm* invasins belong to the internalin protein family and mediate protein-protein interactions through characteristic N-terminal leucin-rich repeats. For example, internalin A (InlA) and internalin B (InlB) interact with and activate their respective host plasma membrane receptors, E-cadherin and c-Met, resulting in phagocytosis of *Lm* by a variety of cell types (3, 4). In addition to epithelial cells and hepatocytes, *Lm* invades and survives in professional phagocytes. These cells not only serve as a niche for *Lm* survival and replication but also facilitate transmission to internal tissues (5). *Lm* uptake by phagocytic cells is thought to occur through C3bi and C1q complement receptors and phagocyte scavenger receptors (6, 7). However, alternative mechanisms of phagocytic cell invasion have not been characterized in detail.

Recently, we reported that interferon-inducible receptor FcγRIa promotes *Lm* invasion of human monocytic cells (8). FcγRIa (also known as CD64) is a high-affinity immunoglobulin G receptor (IgG) receptor, expressed on the surface of monocytes, macrophages, and activated neutrophils (9). FcγRIa itself lacks any known signaling motifs, and upon binding to IgG-opsonized pathogens interacts with the accessory immunoreceptor tyrosine-based activation motif (ITAM)-containing γ-chain (FcεRIg). The γ-chain is then phosphorylated and serves as docking site for intracellular signaling molecules. This allows for FcγRIa-induced downstream events, including pathogen uptake and destruction as well as production and release of inflammatory mediators (10). Strikingly, we previously found that *Lm* is internalized by FcγRIa-expressing cells independently of antibody interaction or ITAM-mediated signaling, hallmarks of IgG-opsonized particles uptake. Additionally, this process did not involve well-characterized host *Lm* internalization receptors (E-cadherin and c-Met) or bacterial surface internalins (InlA and InlB) (8). Thus, FcγRIa-mediated *Lm* entry occurs through a non-canonical mechanism. Interestingly, while FcγRIa enhanced internalization of pathogenic *Lm*, we did not observe increased infection rates of FcγRIa-expressing cells with other intracellular pathogens, including *Shigella flexneri* and *Salmonella* Typhimurium (8), suggesting that a *Lm*-specific factor was involved. However, the identity of this putative FcγRIa-interacting bacterial protein remained unknown.

Here, we set out to identify molecular determinants governing *Lm*-FcγRIa interaction. Predicting that the bacterial ligand of FcγRIa would be a *Listeria* specific virulence factor, we found that FcγRIa does not mediate entry of a nonpathogenic *L. innocua* but enhances internalization of a ruminant pathogen *L. ivanovii*. Consistent with this observation, FcγRIa-mediated *Lm* entry required expression of PrfA, the major activator of *Listeria* virulence. We further identified the PrfA-dependent actin assembly-inducing protein ActA as necessary for FcγRIa-mediated *Lm* entry. Moreover, ActA was sufficient to promote particle uptake by FcγRIa-expressing cells, suggesting that it directly engages FcγRIa on the cell surface. These findings support an alternative *Lm* uptake mechanism by FcγRIa and highlight the role of ActA in host cell invasion.

## Results

### FcγRIa-mediated Lm internalization is blocked by human Fc protein

We previously established that ectopic expression of *FCGR1A* in HEK293A cells was sufficient to reconstitute FcγRIa-mediated *Lm* internalization independently of E-cadherin and c-Met expression, as well as immunoglobulin opsonization of bacteria (8). Here, we employed this simplified system to investigate the molecular interplay involved in *Lm* cellular invasion through FcγRIa. We hypothesized that *Lm* directly engages FcγRIa to trigger internalization. To test this possibility, control and FcγRIa-expressing HEK293A cells were pretreated with excess amounts of recombinant Fc protein (referred to as Fc block) to occupy available surface FcγRIa and prevent interaction with *Lm*. A short-term infection protocol was used to specifically assess *Lm* internalization (11). *Lm* was first incubated with the cells for 1.5 hours to allow initial invasion, and then media was supplemented with gentamicin for 1 hour to kill non-internalized bacteria. Infected cells were lysed to release and quantify colony forming units (CFU) of internalized *Lm*. As shown in Fig. 1A, Fc block did not significantly affect *Lm* uptake by the canonical c-Met-mediated internalization pathway in control cells. However, *Lm* internalization by the FcγRIa pathway was severely abrogated by Fc block (Fig. 1A), indicating that direct bacteria-receptor interaction is indeed required for *Lm* uptake by FcγRIa.

**Figure 1:**
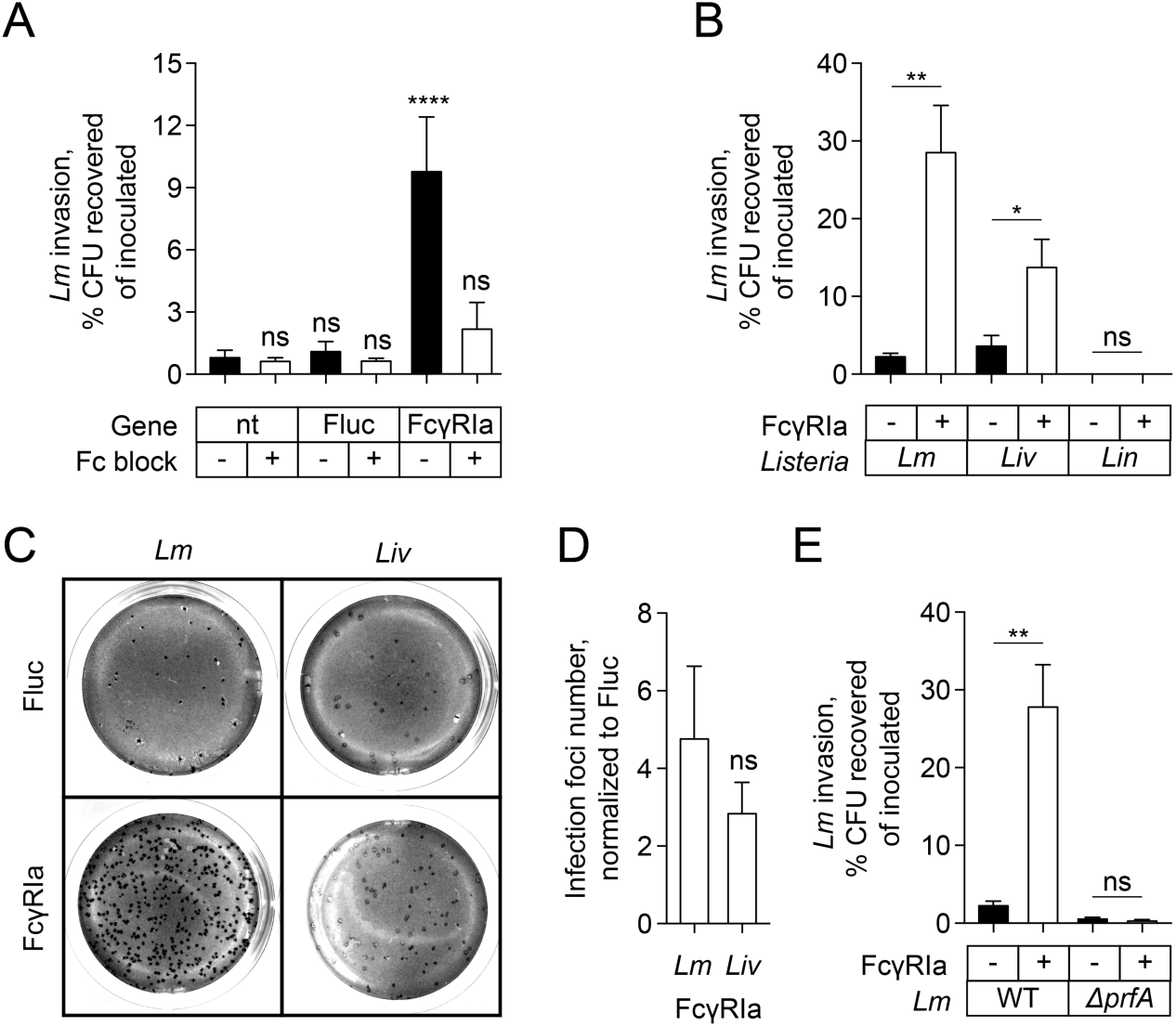
*Listeria* PrfA-dependent virulence is required for FcγRIa-mediated uptake. (A) Invasion of wild type *Lm* in HEK293A cells untransduced or transduced with lentivirus expressing Firefly luciferase (Fluc) or FcγRIa, untreated (black bars) or treated with 0.0125 μg/ml Fc block (white bars). Error bars represent standard deviation (s.d.), n=3. (B) Invasion of wild type *Lm, L.ivanovii* (*Liv*), and *L. innocua* (*Lin*) in HEK293A cells transduced with lentivirus expressing Fluc (black bars) or FcγRIa (white bars). Error bars represent s.d., n=3. (C) Confluent monolayers of HEK293A cells transduced with lentivirus expressing Fluc (*upper*) and FcγRIa (*lower*) and infected with wild type *Lm* (*Lm, left*) and *L. ivanovii* (*Liv, right*). Macroscopic foci were visualized using (3-(4,5-dimethylthiazolyn-2)-2,5-diphenyltetrazolium bromide (tetrazolium MTT) 35 h after initial infection. MOI=0.25 for *Lm*, MOI=0.01 for *L. ivanovii.* (D) Increase in the number of macroscopic infection foci formed by wild type *Lm* (*Lm*) *and L. ivanovii* (*Liv*) in HEK293A cells transduced with lentivirus expressing FcγRIa as compared to HEK293A cells expressing Fluc. Error bars represent s.d., n=4. MOI=0.5, 0.25, 0.05 for *Lm*, MOI=0.05, 0.01, and 0.005 for *L. ivanovii*. (E) Invasion of wild type *Lm* and *Lm ΔprfA* in HEK293A cells transduced with lentivirus expressing Fluc (black bars) or FcγRIa (white bars). Error bars represent s.d., n=3.

### FcγRIa-mediated internalization is specific to pathogenic Listeria species

Our previous studies demonstrated that FcγRIa did not promote cellular uptake of intracellular bacteria *S. flexneri* or *S.* Typhimurium. Additionally, FcγRIa did not confer an invasive phenotype to *L. innocua,* a non-pathogenic *Listeria* species closely related to *Lm* (8). These findings suggest that a bacterial Fcγ receptor ligand is specific to pathogenic *Listeria* and potentially provides an advantage in survival and dissemination during infection. To test this hypothesis, we assessed FcγRIa-mediated internalization of *L. ivanovii*, the only other pathogenic *Listeria* species. This bacterium encodes a *Listeria* pathogenicity island 1 (LIPI-1) with 75% DNA similarity to LIPI-1 found in *Lm* and undergoes the same intracellular lifecycle stages (12). However, unlike *Lm*, *L. ivanovii* is primarily known to cause infection in ruminants with only several human cases reported to date (13-15). Consistent with previous studies, FcγRIa increased cellular invasion of *Lm* 12.7-fold but failed to confer invasiveness to non-pathogenic *L. innocua* (Fig. 1B). Importantly, internalization of *L. ivanovii* was reproducibly potentiated 3.8-fold in presence of FcγRIa as compared to control luciferase-expressing cells (Fig. 1B). This phenotype was further confirmed by visualizing infection foci formed by invading *Lm* and *L. ivanovii* in HEK293A cell monolayers. As expected, the number of *L. ivanovii* infection foci was significantly increased in presence of FcγRIa, similar to that of *Lm* (Figs 1C and 1D). Together, these findings corroborate our initial hypothesis that the FcγRIa-interacting bacterial protein is shared by pathogenic *Listeria* and is not expressed by related non-pathogenic species.

In *Listeria*, transition from a saprotrophic free-living bacterium to an intracellular pathogen is mediated by the transcriptional activator positive regulatory factor A (PrfA) that controls virulence gene expression. Consistent with its role in virulence, PrfA is present in both *Lm* and *L. ivanovii* but not *L. innocua* (16, 17). We next asked whether FcγRIa-mediated *Lm* uptake required expression of PrfA-dependent proteins. The *prfA*-deficient *Lm* strain exhibited decreased internalization in control luciferase-expressing cells (Fig. 1E), likely due to the reduced expression of InlA and InlB (18). Importantly, FcγRIa failed to potentiate uptake of this mutant *Lm* (Fig. 1E), suggesting that expression of the bacterial FcγRIa*-*interacting protein is encoded by a PrfA-regulated gene.

### ActA is necessary for Lm uptake by FcγRIa

PrfA induces expression of approximately 70 genes, with a core set of 12 positively regulated genes, each proceeded by a PrfA box (19). This core group includes PrfA itself and the major virulence factors InlA, InlB required for host cell invasion, LLO, PlcA, and PlcB responsible for vacuolar escape of *Listeria*, metalloprotease Mpl involved in phospholipase maturation, secreted InlC implicated in protrusion formation during bacterial cell-to-cell spread, hexose phosphate transporter Hpt required for uptake of phosphorylated carbohydrates within host cytoplasm, and actin-assembly inducing protein ActA. Finally, products of PrfA-upregulated genes *lmo2218* and *lmo0788* have not been characterized (19-27) (Fig. 2A). We predicted that the bacterial ligand of FcγRIa would be associated with *Lm* surface to facilitate interaction with host cells. As shown in Fig. 2A, only three of the candidate PrfA-upregulated genes are associated with the *Lm* surface via a single anchoring domain (InlA, InlB, ActA). Therefore, we first tested if FcγRIa expression increased invasion levels of *Lm* lacking these genes. Consistent with our prior observations (8), *Lm ΔinlAΔinlB* exhibited increased internalization in presence of FcγRIa, indicating that InlA and InlB were not required for FcγRIa-mediated *Lm* invasion (Fig. 2B). In contrast, uptake of *Lm ΔactA* was not increased by FcγRIa (Fig. 2B), suggesting potential involvement of ActA in FcγRIa-mediated *Lm* invasion. This finding is additionally strengthened by the observation that three other core PrfA-dependent proteins LLO, PlcA and PlcB were dispensable for *Lm* internalization via FcγRIa (Fig. 2C).

**Figure 2:**
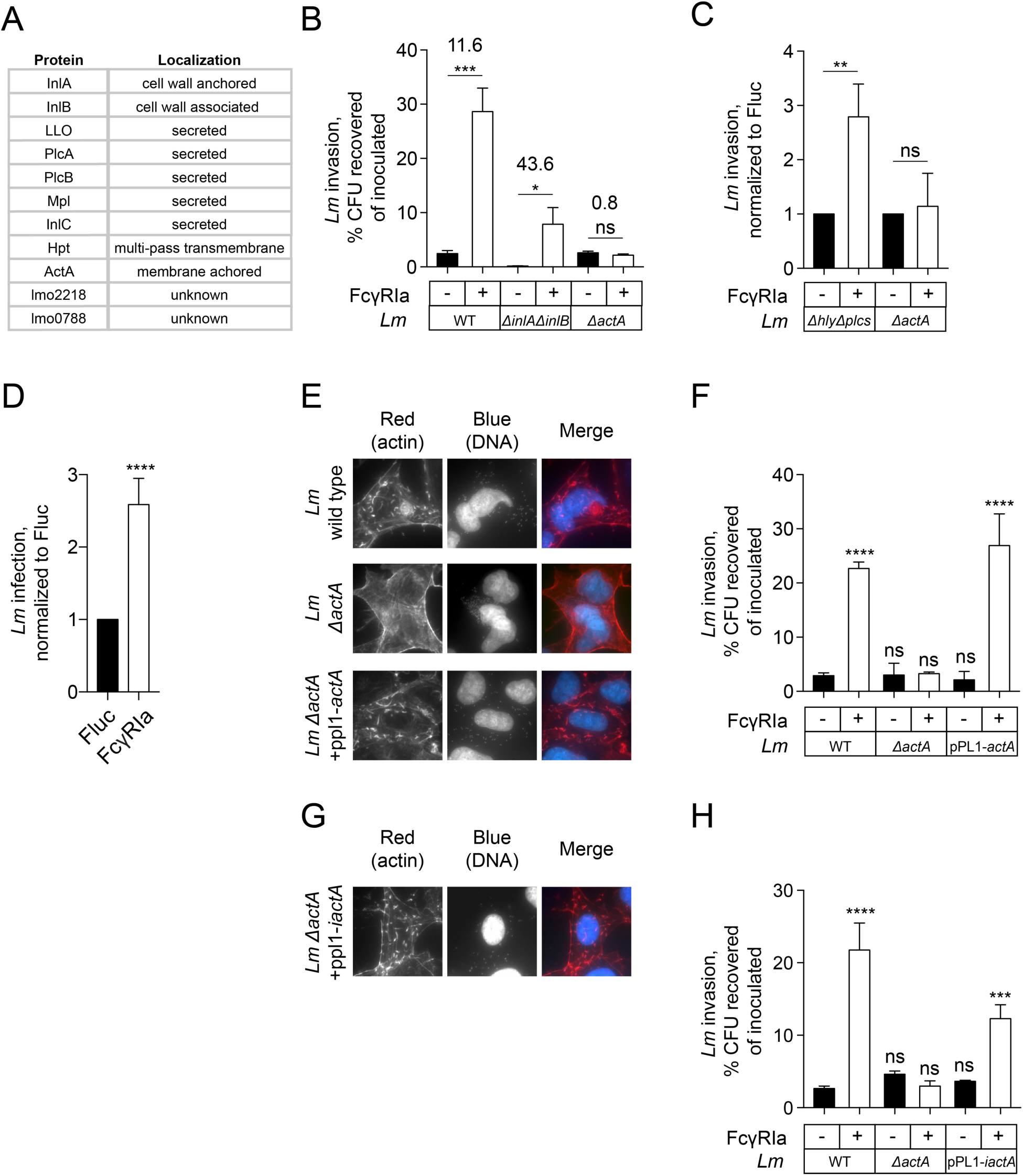
ActA is necessary for FcγRIa-mediated *Lm* invasion. (A) Nomenclature and localization of proteins encoded by the core PrfA-upregulated genes. (B) Invasion of wild type *Lm, Lm ΔinlAΔinlB,* and *Lm ΔactA* in HEK293A cells transduced with lentivirus expressing Fluc (black bars) or FcγRIa (white bars). Error bars represent s.d., n=3. MOI=5. Numbers above bars indicate fold increase in invasion of each strain in HEK293A cells transduced with lentivirus expressing FcγRIa as compared to HEK293A cells expressing Fluc. (C) Invasion of *Lm ΔhlyΔplcAΔplcB* (*ΔhlyΔplcs*) and *Lm ΔactA* in HEK293A cells transduced with lentivirus expressing FcγRIa as compared to HEK293A cells expressing Fluc. Error bars represent s.d., n=3. MOI=5. (D) Infectivity of wild type *Lm* in *CDH1/MET*-deficient HEK293A (clone P4E4) cells transduced with lentivirus expressing Fluc or FcγRIa treated with 1 μM cytochalasin D (see Experimental procedures). Infectivity was quantified as a number of infected GFP-positive cells per well and normalized to Fluc control. Error bars represent s.d., n=6. (E) Fluorescence microscopy of HEK293A cells infected with wild type *Lm, Lm ΔactA,* and *Lm ΔactA* harboring pPL1-*actA* plasmid for 4.5 h following 1.5 h of initial infection. Cells were stained with Alexa Fluor 594 Phallioidin (actin, red) and DAPI (DNA, blue). (F) Invasion of wild type *Lm, Lm ΔactA,* and *Lm ΔactA* harboring pPL1-*actA* in HEK293A cells transduced with lentivirus expressing Fluc (black bars) or FcγRIa (white bars). Error bars represent s.d., n=3. (G) Fluorescence microscopy of HEK293A cells infected with *Lm ΔactA* harboring pPL1-*iactA* plasmid for 4.5 h following 1.5 h of initial infection. Cells were stained with Alexa Fluor 594 Phallioidin (actin, red) and DAPI (DNA, blue). (H) Invasion of wild type *Lm, Lm ΔactA,* and *Lm ΔactA* harboring pPL1-*iactA* in HEK293A cells transduced with lentivirus expressing Fluc (black bars) or FcγRIa (white bars). Error bars represent s.d., n=3.

*Lm* surface protein ActA has been primarily characterized as a mediator of F-actin-driven intracellular *Lm* motility, facilitating dissemination of bacteria throughout the infected tissue (28). ActA deficiency renders *Lm* incapable of intracellular motility and intercellular spread and results in microcolony formation in initially infected cells (29). One potential explanation of the requirement for ActA would be that FcγRIa acts on cell-to-cell spread and its effect is therefore eliminated in the absence of actin-based motility. Short-term infection assays, which allowed us to quantify *Listeria* invasion before any significant amount of cell-to-cell spread occurred, suggested that FcγRIa mediated an increased infection phenotype exclusively by potentiating primary *Lm* internalization (Fig. 2B). However, to further rule out any potential interference of actin-based motility via ActA, we treated FcγRIa-expressing and control cells with actin-destabilization drug cytochalasin D 1 hour following initial *Lm* invasion. While bacterial cell-to-cell spread was blocked, FcγRIa-mediated increase in *Lm* infection was preserved, confirming our prior observations and suggesting a novel actin-independent function of ActA (Fig. 2D).

To verify requirement for ActA in *Lm* invasion, we complemented *Lm ΔactA* with a single copy of *actA* with its proximal promoter (30). Expression of *actA* rescued actin-mediated motility in a cell-to-cell spread deficient *Lm ΔactA* as observed by actin comet tail formation (Fig. 2E). Importantly, introduction of *actA* also restored FcγRIa-mediated internalization to the level of wild type *Lm* (Fig. 2F), confirming requirement of ActA for *Lm* uptake by FcγRIa. FcγRIa also increased invasion of *L. ivanovii* (Fig. 1B), which expresses an actin-assembly inducing protein iActA. *L. ivanovii* iActA and *Lm* ActA share only 34% amino acid identity, however, the overall protein architecture is conserved along with several regions of close homology (31, 32). Further, we complemented *Lm ΔactA* with *iactA* and its proximal promoter. Notably, both actin-based motility and FcγRIa-mediated internalization were restored with introduction of iActA (Figs 2G and 2H). Taken together, these results suggest that ActA is a key protein involved in FcγRIa-mediated *Listeria* uptake.

### ActA is sufficient for Lm uptake by FcγRIa

We next sought to determine if ActA was sufficient to promote *Lm* entry via FcγRIa or whether other bacterial proteins present in both *Lm* and *L. ivanovii* were involved. We used passive absorption to coat 1-μm-diameter carboxylated polystyrene beads with purified His-tagged extracellular domain of ActA and measured their uptake by FcγRIa-expressing HEK293A cells. Similar approaches have been successfully used to study *Lm* invasion, for example, to confirm the role of internalin A in *Lm* internalization by E-cadherin expressing cells (33). In addition, we previously reconstituted human IgG-coated bead uptake by FcγRIa in nonprofessional phagocytic cells (8). Here, polystyrene beads were preincubated with ActA-His and MBP-His as a control for 1 hour and washed in BSA-containing blocking buffer. As shown in Fig. 3A, both proteins were efficiently absorbed by the beads. ActA- or MBP-His-coated beads were then incubated with FcγRIa or luciferase-expressing cells at 37°C to allow bead binding and internalization. After 1.5 hours, cells were placed on ice to inhibit bead entry and stained with anti-His tag antibodies without permeabilization to distinguish between cell-associated and internalized beads (Fig. 3B). While only low background levels of internalization were observed for either type of beads in luciferase-expressing cells, FcγRIa facilitated uptake of ActA-coated but not MBP-coated beads (35.33±1.606% compared to 5.746±0.616%, n=3 independent replicates) (Figs 3C and 3D). Further, specificity of the uptake was confirmed by incubating cells with Fc block prior to addition of ActA-His-coated beads to occupy available surface FcγRIa. Similar to the entry of ActA-expressing *Lm* (Fig 1A), Fc block treatment inhibited FcγRIa-mediated uptake of Act-His-coated beads (Fig 3E). Collectively, these data indicate that ActA is sufficient to induce FcγRIa-dependent internalization.

**Figure 3:**
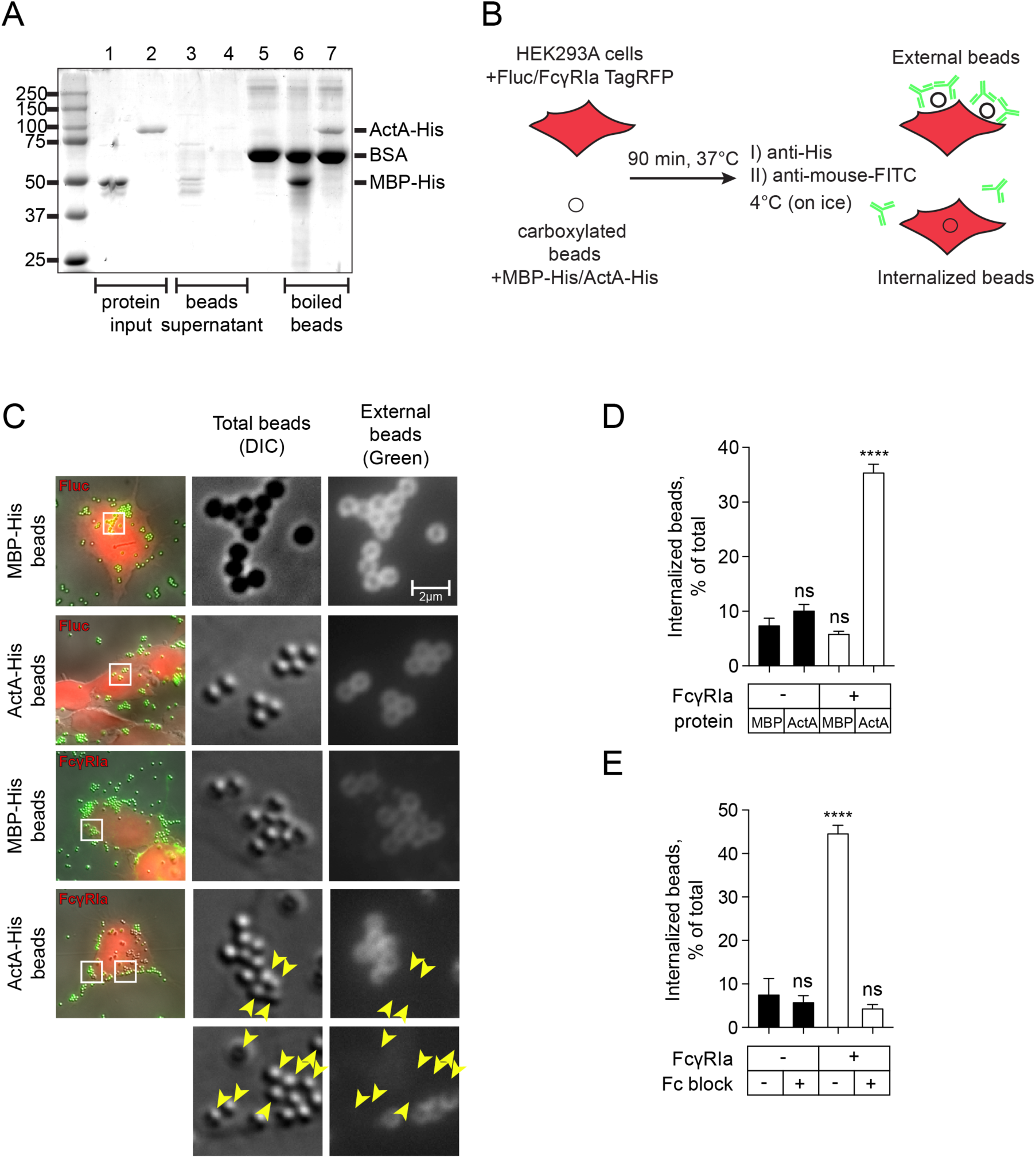
ActA is sufficient to confer invasiveness to polystyrene beads. (A) Proteins analyzed by SDS-PAGE and stained with Coomassie Blue R-250. *Lane 1:* MBP-His input used for bead coating (10%), *Lane 2:* ActA-His input used for bead coating (10%), *Lane 3:* supernatant of MBP-His-coated beads (unabsorbed protein, 20%), *Lane 4:* Supernatant of ActA-His-coated beads (unabsorbed protein, 20%), *Lane 5:* PolyLink Wash/Storage Buffer [10 mM Tris (pH 8.0), 0.05% BSA, 0.05% Proclin 300], *Lane 6:* MBP-His absorbed on beads (20%), *Lane 7:* ActA-His absorbed on beads (20%). Precision Plus Protein(tm) Prestained Standards (Bio-Rad) was used for molecular weight estimation (left lane). (B) Diagram illustrating the phagocytic assay used to reconstitute FcγRIa-mediated uptake of MBP-His or ActA-His-coated beads (C) Fluorescence and differential interference contrast (DIC) microscopy of HEK293A cells transduced with lentivirus expressing Fluc or FcγRIa and incubated with MBP-His or ActA-His-coated beads. Arrows indicate internalized beads. (D) Quantification of internalized MBP-His and ActA-His-coated beads in HEK293A cells transduced with lentivirus expressing Fluc (black bars) or FcγRIa (white bars). Error bars represent s.d., 15 cells were counted for each of three independent experiments. (E) Quantification of internalized ActA-His-coated beads in HEK293A cells transduced with lentivirus expressing Fluc (black bars) or FcγRIa (white bars), mock-treated or treated with 0.0125 μg/ml Fc block prior to the assay. Error bars represent s.d., 15 cells were counted for each of three independent experiments.

### Actin nucleation ability of ActA is dispensable for Lm invasion via FcγRIa

ActA has recently emerged as a multifunctional virulence factor, implicated in epithelial cell invasion, phagosome escape, and autophagy evasion in addition to its well-established role in actin-based motility and bacterial cell-to-cell spread (28, 34). To distinguish between ActA function in cytoplasmic motility and its newly identified role in FcγRIa-mediated entry, we asked whether *Lm* strains expressing *actA* mutants with abrogated actin nucleating activity preserved their ability for FcγRIa-dependent invasion. ActA encodes a cofilin homology region between residues 135 and 165 that is homologous to the proteins of the Wiscott-Aldrich Syndrome protein (WASP) family and is critical for actin-based motility (35). Within the cofilin region, the ^146^KKRRK^150^ motif is essential for recruitment of actin-nucleation complex Arp2/3, allowing ActA to serve as an actin nucleation-promoting factor (36-39) (Fig. 4A). As expected, *Lm* expressing a minimal (Δ146-150) as well as a full deletion of the cofilin homology region (Δ135-165) did not form actin comet tails and failed to disseminate in the monolayer of HEK293A cells (Fig. 4B). However, these strains readily invaded FcγRIa-expressing cells, with an increase in invasion of 9.23 and 8.49-fold, respectively, compared to luciferase-expressing cells (Fig. 4C). We concluded that ActA function in FcγRIa-mediated invasion is independent of its role in actin nucleation and cell-to-cell spread. Additionally, cofilin homology region (135-165 a.a.) is not involved in ActA-FcγRIa interaction.

**Figure 4:**
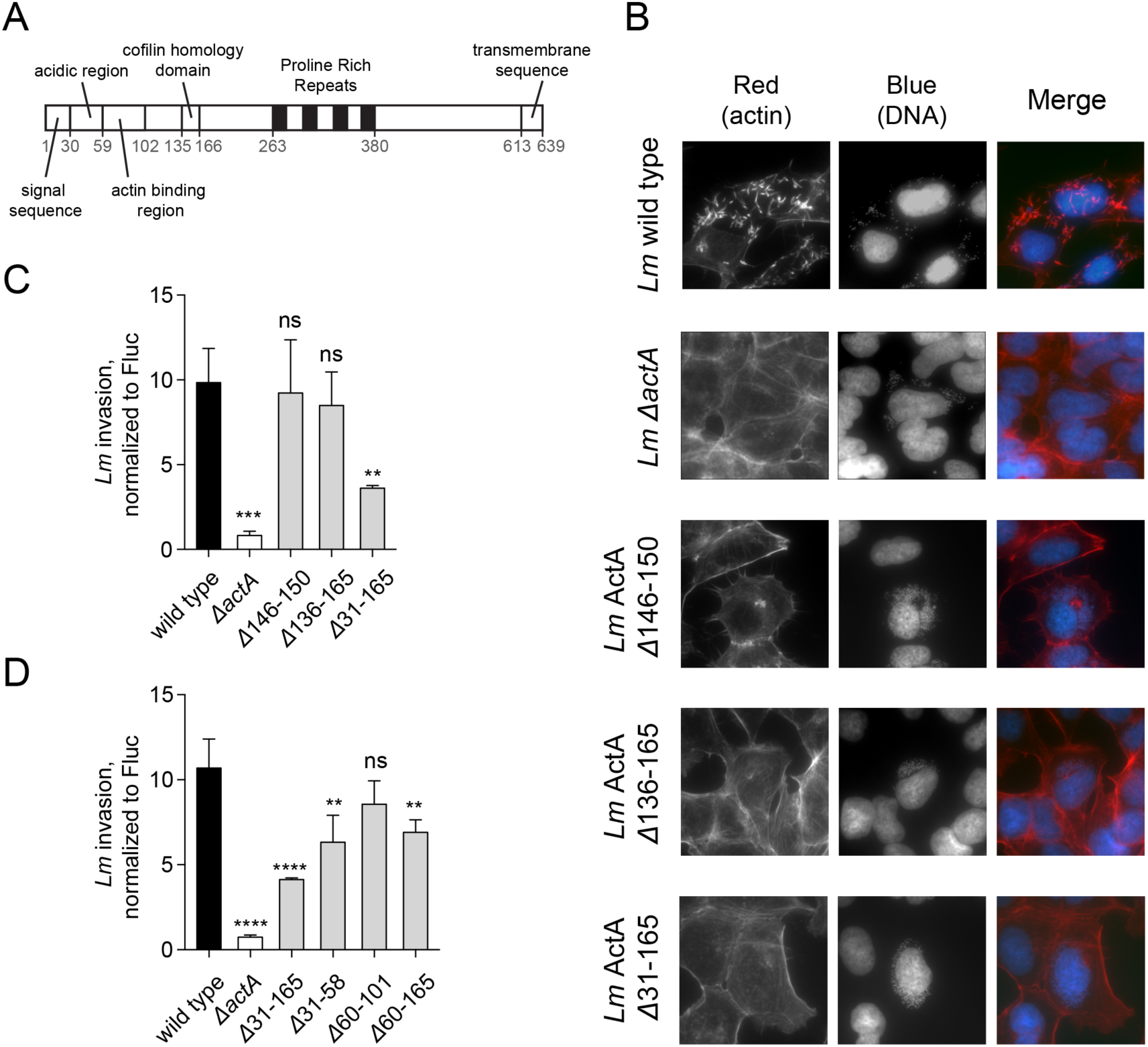
FcγRIa-mediated *Lm* internalization is independent of the ActA-induced actin polymerization. (A) Schematic diagram of *Lm* ActA domain structure. (B) Fluorescence microscopy of HEK293A cells infected with *Lm* ActA Δ146-150, *Lm* ActA Δ136-165, *Lm* ActA Δ31-165 for 4.5 h following 1.5 h of initial infection. Cells were stained with Alexa Fluor 594 Phallioidin (actin, red) and DAPI (DNA, blue). (C) Fold increase in wild type *Lm, Lm ΔactA*, *Lm* ActA Δ146-150, *Lm* ActA Δ136-165, and *Lm* ActA Δ31-165 invasion of HEK293A cells transduced with FcγRIa-expressing lentivirus as compared to HEK293A cells expressing Fluc. Error bars represent s.d., n=3. (D) Fold increase in wild type *Lm, Lm ΔactA*, *Lm* ActA Δ31-165, *Lm* ActA Δ31-58, *Lm* ActA Δ60-101, and *Lm* ActA Δ60-165 invasion of HEK293A cells transduced with FcγRIa-expressing lentivirus as compared to HEK293A cells expressing Fluc. Error bars represent s.d., n=3.

Lastly, to determine if the N-terminal region of ActA (31-165 a.a.) is involved in FcγRIa-mediated uptake, we infected FcγRIa-expressing cells with *Lm* expressing ActA Δ31-165. Interestingly, invasion of this mutant strain was significantly attenuated compared to wild type *Lm* (Fig. 4C). Since deletion of the cofilin homology domain (135-165 a.a.) did not affect internalization, we concluded that the 31-135 a.a. region is important for the FcγRIa-mediated uptake. To identify the minimal region of the ActA N-terminal domain required for FcγRIa-dependent internalization, we tested *Lm* invasiveness conferred by ActA deletion variants Δ31-58, Δ60-101, and Δ60-165. While the largest deletion Δ31-165 exhibited reduced level of internalization in FcγRIa-expressing cells, none of the smaller deletions (Δ31-58, Δ60-101, Δ60-165) alone caused a comparable invasion defect (Fig. 4D). These results suggest that multiple regions within the N-terminal region of ActA (amino acids 31-165), excluding the cofilin homology domain are required for FcγRIa-mediated internalization.

## Discussion

We previously identified FcγRIa as an interferon-stimulated gene involved in *Lm* uptake by myeloid cells (8). FcγRIa-mediated uptake of *Lm* proceeded through a novel mechanism, independent of IgG opsonization or canonical signaling through the immunoreceptor tyrosine-based activation motif (ITAM). However, it remained unclear how *Lm* recognized FcγRIa for internalization. Here, we establish ActA, an actin assembly-inducing protein, as a bacterial surface ligand employed by *Lm* to promote uptake by FcγRIa-expressing cells. Our genetic and biochemical experiments provide compelling evidence that ActA is both necessary and sufficient for FcγRIa-dependent *Lm* internalization independently of its actin-nucleation function.

Suspecting a role of FcγRIa-mediated *Lm* uptake in *Lm* virulence, we narrowed down the search for a bacterial determinant of *Lm*-FcγRIa interaction to PrfA-activated genes. Among those, *actA* was identified as the primary candidate for *Lm* internalization by FcγRIa. ActA was first described in 1992 as a PrfA-regulated gene product, required for *Lm*-induced actin assembly and cellular dissemination (26, 29). Since then the mechanism of *Lm* actin-based motility has been characterized in detail (37, 40, 41). However, a growing number of reports indicate that in addition to its canonical role in cytosolic motility and cell-to-cell spread, ActA contributes to other stages of *Lm* infection (28). For example, it was found to be involved in *Lm* adhesion and invasion of various cell types, potentially though interacting with heparan sulfate proteoglycans (HSPG) on the host cell surface (42, 43). Further, it was shown to participate in *Lm* escape from autophagic recognition, while initial evidence also suggested its role in promoting maturation of *Lm*-containing phagosomes (44-46). Lastly, ActA mediates bacterial aggregation and biofilm formation, ultimately facilitating host colonization (47). Despite its clear role in multiple host cell processes, the molecular interaction between ActA and host substrates that facilitate these processes remain to be characterized. Thus, this study implicating ActA in FcγRIa-mediated internalization sheds light on this multifunctional virulence factor.

We have previously found that FcγRIa selectively promotes internalization of pathogenic *Lm* but not other intracellular bacteria including *S.* Typhimurium and *S. flexneri* or nonpathogenic *Listeria* species (8). Identification of ActA as a protein required for *Lm* uptake via FcγRIa provides an explanation for the observed bacterial species specificity. ActA is a protein unique to *Listeria* pathogens and its homologues are not found in such intracellular bacteria such as *S.* Typhimurium. Interestingly, like *Lm*, *S. flexneri* encodes the actin-nucleating factor IcsA that promotes cell-to-cell spread. However, ActA and IcsA are not homologous and use distinct mechanisms for actin polymerization (48). ActA is also absent from non-pathogenic *L. innocua*, a close relative of a highly pathogenic *Lm* (17). The only close homologue of *Lm* ActA is an actin nucleating protein iActA expressed by a ruminant pathogen *L. ivanovii* (31, 49). Additionally, homologous protein sActA is encoded but not expressed by nonpathogenic *L. seeligeri* (50). Consistent with the role of ActA in *Listeria* uptake, FcγRIa expression increase internalization of *L. ivanovii* (as reported here) but did not facilitate uptake of *S. flexneri* (8). *Lm* and *L. ivanovii* actin assembly-inducing proteins only share 34% amino acid identity, however, their protein architecture and several functional regions are highly conserved (31, 32). While we have determined that the cofilin homology domain is not required for *Lm* invasion, other conserved domains of ActA, such as proline-rich repeats might be involved.

One potential complication of our model is that ActA expression was originally believed to be restricted to the host intracellular environment due to its regulation by PrfA (51, 52). However, its expression is controlled by two distinct promoters, only one of which is PrfA-dependent (53). Accumulating evidence suggests that ActA is expressed in the gut lumen during initial adhesion, prior to invasion (47, 54). Therefore, it is possible that under *in vivo* conditions ActA would be expressed by extracellular bacteria, promoting internalization via FcγRIa-expressing myeloid cells.

A key question that emerges from our work is how ActA contributes to *Lm* internalization by FcγRIa. Here, we uncovered ActA as necessary and sufficient for FcγRIa-mediated *Lm* internalization, however, whether these proteins serve as the primary receptor pair remains unclear. Our previous study demonstrated that transmembrane and intracellular domains of FcγRIa were dispensable for receptor-mediated *Lm* internalization (8). Therefore, we predict that a co-receptor initiating downstream signaling and FcγRIa uptake is likely to be involved. Our current data suggest direct interaction of ActA and FcγRIa, since ActA-coated polystyrene beads were internalized similarly to ActA-expressing bacteria. It is also possible that a multiprotein interaction involving a co-receptor is occurring. Thus, in the future, it will be critical to determine whether such accessory protein is indeed involved and if ActA directly binds FcγRIa and/or the co-receptor. In addition, it will be important to pinpoint the downstream signaling events, mediating *Lm-*FcγRIa internalization.

In summary, our results uncover an unexpected new role of ActA in *Lm* invasion through a human immunoglobulin receptor FcγRIa. This study strengthens our understanding of FcγRIa-mediated *Lm* uptake mechanism and adds to a growing number of ActA contributions to a successful *Lm* lifecycle.

## Acknowledgements

The authors would like to specifically thank Dan Portnoy (UC Berkeley) for helpful suggestions and providing *Listeria* strains; and all members of the Alto laboratory for support and valuable discussions.

## Funding

This research was supported by grants to N.M.A. from the National Institute of Health (AI083359), the Welch Foundation (#I-1831), The Burroughs Welcome Fund, and the Howard Hughes Medical Institute and Simons Foundation Faculty Scholars Program. M.E.A was supported by the UTSW Molecular Microbiology Training Grant (49C0035803).

## Author contributions

S.S.P., M.E.A., and N.M.A. designed the study, developed the methodology, and wrote the manuscript. S.S.P. and M.E.A. collected the data and performed the analysis.

## Experimental procedures

### Bacterial culture

Bacterial strains used in this study are listed in Table 1. All *Listeria* strains were maintained in Brain Heart Infusion (BHI) media (Difco, BD Biosciences) with appropriate antibiotics (see Table 1). All Lm strains were streptomycin resistant (up to 200 μg/ml). *Escherichia coli* strains (DH5 alpha, SM10, and BL21(DE3)) were maintained in Luria-Bertani (LB) media with appropriate antibiotics (see Table 1).

**Table 1.** Bacterial strains used in this study

| Strain | Description | Source |
| --- | --- | --- |
| *Escherichia coli* DH5α | *E. coli* strain used for general cloning procedures | Thermo Fisher Scientific |
| *E. coli* BL21(DE3) | *E. coli* strain used for use with bacteriophage T7 promoter-based expression systems | Thermo Fisher Scientific |
| *E. coli* SM10 λpir | Used as a donor strain for bacterial conjugation, kanamycin resistant (50 µg/ml) | Provided by Sebastian Winter (UTSW) |
| *Listeria monocytogenes* 10403s, GFP | Wild type *L. monocytogenes* 10403s strain with constitutive GFP expression | Provided by Dan Portnoy (UC Berkley) |
| *L. monocytogenes* DP-L3078, pPL2-pactA::GFP | *L. monocytogenes* Δ*actA* with pactA-dependent GFP expression, chloramphenicol resistant (25 µg/ml) | (8) |
| *L. monocytogenes* DP-L2319, pPL2-pactA::GFP | *L. monocytogenes* Δ*hly* Δ*plcA* Δ*plcB* with pactA-dependent GFP expression, chloramphenicol resistant (25 µg/ml) | (8) |
| *L. monocytogenes* DP-L4317 | *L. monocytogenes* Δ*prfA* | Provided by Dan Portnoy (UC Berkley) (57) |
| *L. monocytogenes* DP-L3992 | *L. monocytogenes* expressing ActA Δ146-150 | Provided by Dan Portnoy (UC Berkley) (35) |
| *L. monocytogenes* DP-L3990 | *L. monocytogenes* expressing ActA Δ136-165 | Provided by Dan Portnoy (UC Berkley) (35) |
| *L. monocytogenes* DP-L3984 | *L. monocytogenes* expressing ActA Δ31-165 | Provided by Dan Portnoy (UC Berkley) (35) |
| *L. monocytogenes* DP-L3996 | *L. monocytogenes* expressing ActA Δ31-58 | Provided by Dan Portnoy (UC Berkley) (35) |
| *L. monocytogenes* DP-L3994 | *L. monocytogenes* expressing ActA Δ60-101 | Provided by Dan Portnoy (UC Berkley) (35) |
| *L. monocytogenes* DP-L3986 | *L. monocytogenes* expressing ActA Δ60-165 | Provided by Dan Portnoy (UC Berkley) (35) |
| *L. monocytogenes* DP-L4029 | Phage-cured *L. monocytogenes* Δ*actA* | Provided by Dan Portnoy (UC Berkley) (30) |
| *L. ivanovii* subsp. *ivanovii* Seeliger et al. (DP-L393) | Wild type *L. ivanovii* strain | Provided by Dan Portnoy (UC Berkley) (ATCC 19119) |
| *L. innocua* Seeliger CLIP 11262 | Genome sequencing *L. innocua* strain | ATCC BAA-680 |
| *L. monocytogenes* DP-L4029 + pPL1-*actA* | DP-L4029 complemented with *L. monocytogenes* ActA, chloramphenicol resistant (25 µg/ml) | This study |
| *L. monocytogenes* DP-L4029 + pPL1-*iactA* | DP-L4029 complemented with *L. ivanovii* iActA, chloramphenicol resistant (25 µg/ml) | This study |

### Mammalian cell culture

HEK293A (kindly provided by Jack Dixon, UC San Diego) and *CDH1/MET*-deficient HEK293A (clone P4E4) cells (previously described in (8)) were maintained in Dulbecco’s Modified Eagle Medium (DMEM) (Gibco), supplemented with 10% Fetal Bovine Serum (FBS) (Gibco) and non-essential amino acids (NEAA) (Gibco).

### DNA constructs

Lentiviral vector constructs pTRIP.CMV.IVSb.ires.TagRFP-Fluc (Firefly luciferase) and pTRIP.CMV.IVSb.ires.TagRFP-FCGR1A were provided by Dr. John Schoggins (UTSW).

pPL1-*actA* and pPL1*-iactA* were generated as previously described (30). Briefly, *actA* gene and its proximal promoter were cloned into pPL1 using primers described in (30) with BamHI and NotI restriction sites:

actA_F_Bam: atgcGGATCCtgaagcttgggaagcag, actA_R_Not: atgcGCGGCCGCtcaagcacatacctag.

Similar procedure was used to clone *iactA* of *L. ivanovii* with the following primers: iactA_F_Bam: atgcGGATCCatacaggcggaagaagtgg, iactA_R_Not: atgcGCGGCCGCtttgttgattctcatagcattatgt.

pET28-6xHis-MBP-6xHis was provided by the Alto laboratory.

### Generation of lentiviral pseudoparticles

Lentiviral pseudoparticles were generated as previously described (55).

### Lentiviral transduction

Lentiviral transduction was performed as previously described (8). Briefly, cells were seeded in 24-well tissue culture plates at a density of 7x10^4^ cells per well and transduced the following day with lentiviral pseudoparticles via spinoculation at 1,000 x *g* for 45 min in medium containing 3% FBS, 20mM HEPES and 4 μg/ml polybrene. 6 h after spinoculation, pseudoparticle-containing media was removed and replaced with full cell culture medium, containing 10% FBS and NEAA. For subsequent assays, cells were split 48 h - 72 h after transduction.

### Bacterial conjugation

Conjugation was used to introduce pPL1 integration vectors into *L. monocytogenes*. Briefly, pPL1 derived plasmid constructs were first chemically transformed into *E. coli* strain SM10 (56) by using standard procedures. Transformed SM10 (donor) were grown at 30°C with shaking to mid-log phase (optical density at 600 nm, OD600 ∼ 0.6) in LB medium supplemented with 25 μg/ml of chloramphenicol. Recipient phage-cured *L. monocytogenes* were grown at 30°C with shaking to midlog phase in antibiotic-free BHI medium. Donor culture (250 μl) was harvested by centrifugation and washed twice with antibiotic-free BHI before combining with the recipient culture (150 μl). The mixture was plated on BHI plates and incubated at 30°C for at least 12 h. Following incubation, bacteria were streaked out into 150 μl sterile water, and 20 μl of resulting suspension plated on BHI plates supplemented with 7.5 μg/ml of chloramphenicol and 50 μg/ml of streptomycin. The plates were then incubated at 37°C for 36 h. Individual colonies were picked and screened by PCR for integration with primers PL14 and PL61 (30).

### *Listeria* infection

*Listeria* was inoculated from a frozen stock and grown for 13 h at 30°C in BHI media with appropriate antibiotics without shaking. 1 ml of bacteria was then washed in phosphate buffer saline (PBS) and resuspended in 1 ml of PBS. Bacteria were then added to each well of cells to achieve a multiplicity of infection (MOI) of 5 (unless otherwise stated in the corresponding figure legend) and incubated for 90 min at 37°C, 5% CO_2_. Culture media was then removed and replaced with media supplemented with 25 μg/ml gentamicin (Quality Biological) and cells incubated at 37°C, 5% CO_2_ for 1 h. Following *Listeria* infection, mammalian cells were washed three times with PBS and then lysed by incubating in 0.5% Triton X-100 / PBS for 5 min at room temperature, followed by vigorous pipetting to complete the lysis. Intracellular bacterial burden was determined by plating serial dilutions of suspension on BHI-agar plates, incubating at 37°C overnight, and counting bacterial colony forming units (CFU) the next day. Additionally, serial dilutions of bacterial culture used for infection were plated to obtain the inoculated CFU. Finally, the following equation was used to calculate *Listeria* invasion: % invasion = [CFU recovered per well/CFU inoculated per well] x 100%.

For the Fc block experiment, prior to infection cells were pretreated with 200 μl of 0.0125 μg/ml Fc block (BD Biosciences 564219) in 10% FBS / DMEM or mock-treated with 10% FBS / DMEM for 15 min, and washed once with 10% FBS / DMEM. Bacteria were then added to each well of cells and incubated for 2 h at 37°C, 5% CO_2_. Gentamicin treatment and collection were performed as described above.

To visualize bacterial infection by epifluorescence microscopy, infection was performed in cells cultured in 4-well chamber slides (Falcon). Following infection, cells were washed once in PBS, fixed in 3.7% formaldehyde in PBS for 10 min at room temperature, and permeabilized with 0.5% Triton X-100 / PBS. Actin was visualized by incubating cell with Alexa Fluor 594 Phallioidin (Molecular Probes, Thermo Fisher Scientific). Cells were then washed three times in PBS and incubated for 2 min in 4’,6-diamidino-2-phenylindole (DAPI) solution. Samples were washed 5 times with PBS and chamber removed from the slide. When the excess liquid dried, the coverslips were mounted on the samples with ProLong Gold reagent (Molecular Probes, Life Technologies). Samples were observed with a fluorescent microscope Zeiss Observer Z1.

For cytochalasin treatment experiment, cells were infected with wild type GFP-expressing *Lm* for 1 hour, followed by treatment with 1 μM cytochalasin D (Cayman Chemical) and 25 μg/ml gentamicin. Samples were processed in 24-well plates 6 hours after treatment as described earlier and imaged using Nikon Eclipse Ti Fluorescence Microscope. Total of 64 images were acquired per well, which covered >90% of the well area.

### Agarose overlay assay

Agarose overlay assays were performed as previously described (8).

### Protein purification

*ActA-His:* ActA (1-613)-6xHis was purified as previously described (37). Briefly, 10 ml of BHI media supplemented with 4 μg/ml chloramphenicol was inoculated with DP-L2723 from a frozen stock. Culture was incubated at 37°C for 12 h with shaking. Resulting culture was then used to inoculate 400 ml of BHI media (4 μg/ml chloramphenicol) and grown at 37°C for 12 h with shaking. Bacteria were then pelleted (2,700 x *g* for 20 min) and protein precipitated from the culture supernatant by slowly adding ammonium sulfate to 60% saturation (312 g per 1L). Precipitation was allowed to proceed for at least 2 h at 4°C with mixing. Precipitated protein was pelleted (7000 x *g* for 30 min at 4°C) and resuspended in 20 ml resuspension buffer [20 mM HEPES (pH 7.5), 50 mM KCl, 10 mM imidazole, 1 mM Dithiothreitol (DTT), 1x SIGMAFAST Protease Inhibitor Cocktail Tablets, EDTA-Free] and bound to 1 ml of nickel nitrilotriacetic acid (Ni-NTA) agarose resin (Qiagen) for 1.5 h at 4°C. The resin was loaded on a column, washed with 40 ml of wash buffer [20 mM HEPES (pH 7.5), 50 mM KCl, 25 mM imidazole, 1 mM DTT] and ActA-His eluted with 8 ml of elution buffer [20 mM HEPES (pH 7.5), 50 mM KCl, 500 mM imidazole]. Eluted protein was further concentrated and buffer exchanged 1:2000 into storage buffer [20 mM HEPES (pH 7.5), 50 mM KCl] using Amicon Ultra-15 Centrifugal Filter Unit with Ultracel-30 membrane (Millipore). Typically, a concentration of 200 μg/ml was obtained as determined by BCA Protein Assay Kit (Pierce, Thermo Fisher Scientific). Finally, DTT (to 1mM) and glycerol (to 10%) were added and protein stored at -80°C.

*MBP-His:* pET28-6xHis-MBP-6xHis was chemically transformed into BL21(DE3) cells using standard procedures. 5 ml of LB media supplemented with 50 μg/ml kanamycin was inoculated with BL21(DE3) pET28-His-MBP-His and incubated at 37°C for 12 h with shaking. Resulting culture was then used to inoculate 200 ml of LB media (25 μg/ml kanamycin) and grown at 37°C for 3 h with shaking to OD600 ∼ 0.6. Protein expression was induced by addition of isopropyl β-D-1-thiogalactopyranoside (IPTG) to the media to achieve a final concentration of 0.5 mM. Bacteria were then incubated at 16°C for additional 16 h and pelleted by centrifugation. His-MBP-His (referred to as MBP-His in text) was purified using Amilose Resin (New England Biolabs) according to manufacturer’s instructions. Eluted protein was further concentrated and buffer exchanged 1:2000 into Tris-Buffered Saline (TBS) using Amicon Ultra-15 Centrifugal Filter Unit with Ultracel-10 membrane (Millipore). Finally, DTT (to 1mM) and glycerol (to 10%) were added and protein stored at -80°C.

### *In vitro* bead uptake assay

HEK293A cells were transduced with lentivirus co-expressing TagRFP and Fluc or FcγRIa. 48h or 72h after transduction, cells were plated at 7x10^4^ cells/ml in 4-well chamber slides (Falcon) with poly-lysine coating. The day of the assay 40 μl of carboxylated latex beads (Polysciences, Polybead Carboxylate 1.0 Micron Microspheres, [9003-53-6], 2.7% solids-latex, Cat# 08226, dia = 0.91 μm) were washed twice in PolyLink Coupling Buffer [50 mM MES (pH 5.2), 0.05% Proclin 300] (Polysciences) and preincubated for 10 min with shaking. Beads were then pelleted, resuspended in 100 μl of 200 μg/ml protein solution (ActA-His or MBP-His) and incubated for 1 h at room temperature with constant shaking to allow passive absorption. Unabsorbed proteins were collected from bead supernatant for further analysis by SDS-PAGE (see below). Protein-coated beads were then washed twice in PolyLink Wash/Storage Buffer [10 mM Tris (pH 8.0), 0.05% Bovine Serum Albumin (BSA), 0.05% Proclin 300], resuspended in 1ml 10% FBS / DMEM and 20 μl added to cells in chamber slides. Slides were centrifuged at 800 x *g* for 1 min, and then placed 37°C for 90 min. After the incubation, slides were placed on ice and washed with ice-cold medium to inhibit further bead uptake. Extracellular beads were then labeled with mouse anti-His tag antibody (Aviva Systems Biology, OAEA00010) at 1:250 dilution in 10%FBS/DMEM for 1 h on ice. Cells were washed 5 times with ice-cold PBS and stained with fluorescein isothiocyanate (FITC)-conjugated goat anti-mouse antibody (Thermo Fisher Scientific, 31569) at 1:200 dilution in ice-cold PBS for 30 min. Cells were then fixed with 3.7% PFA for 20 min at room temperature. All samples were washed 5 times with PBS and chamber removed from the slide. When the excess liquid dried, the coverslips were mounted on the samples with ProLong Gold reagent (Molecular Probes, Life Technologies). Samples were observed with a fluorescent microscope Zeiss Observer Z1. The overall number of beads and green extracellular beads was visually evaluated and at least 15 TagRFP-positive cells per sample were used for quantification with three biological replicates per experimental condition. Uptake was measured as a percentage of internalized beads, determined by subtracting the number of extracellular (green) beads from the total beads, divided by the number of total beads.

For the Fc block experiment, prior to infection cells were pretreated with 200 μl of 0.0125 μg/ml Fc block (BD Biosciences 564219) in 10% FBS / DMEM or mock-treated with 10% FBS / DMEM for 15 min, and washed once with 10% FBS / DMEM.

To evaluate protein absorption, beads were resuspended in Laemmli Sample buffer (Bio-Rad) with 2-mercaptoethanol and boiled for 5 min at 95°C. Beads were then pelleted by centrifugation and supernatant analyzed by SDS-PAGE (following standard protocols).

### Statistical analysis

All experiments were performed as three independent replicates, unless otherwise stated. For experiments where only two groups of samples were compared, unpaired t-test was used to determine if difference between groups was statistically significant. To determine statistical significance in experiments with three or more groups of samples, one-way analysis of variance (ANOVA) with Dunnett’s procedure for multiple comparisons was used. Data analysis was performed in GraphPad Prism software.

## References

1. Cossart P, Sansonetti PJ. Bacterial invasion: the paradigms of enteroinvasive pathogens. Science. 2004;304(5668):242–8.

2. Pizarro-Cerda J, Kuhbacher A, Cossart P. Entry of Listeria monocytogenes in mammalian epithelial cells: an updated view. Cold Spring Harbor perspectives in medicine. 2012;2(11).

3. Bierne H, Sabet C, Personnic N, Cossart P. Internalins: a complex family of leucine-rich repeat-containing proteins in Listeria monocytogenes. Microbes Infect. 2007;9(10):1156–66.

4. Pizarro-Cerda J, Charbit A, Enninga J, Lafont F, Cossart P. Manipulation of host membranes by the bacterial pathogens Listeria, Francisella, Shigella and Yersinia. Semin Cell Dev Biol. 2016.

5. Drevets DA, Sawyer RT, Potter TA, Campbell PA. Listeria monocytogenes infects human endothelial cells by two distinct mechanisms. Infection and immunity. 1995;63(11):4268–76.

6. Drevets DA, Campbell PA. Roles of complement and complement receptor type 3 in phagocytosis of Listeria monocytogenes by inflammatory mouse peritoneal macrophages. Infection and immunity. 1991;59(8):2645–52.

7. Alvarez-Dominguez C, Carrasco-Marin E, Leyva-Cobian F. Role of complement component C1q in phagocytosis of Listeria monocytogenes by murine macrophage-like cell lines. Infection and immunity. 1993;61(9):3664–72.

8. Perelman SS, Abrams ME, Eitson JL, Chen D, Jimenez A, Mettlen M, et al. Cell-Based Screen Identifies Human Interferon-Stimulated Regulators of Listeria monocytogenes Infection. PLoS Pathog. 2016;12(12):e1006102.

9. Pincetic A, Bournazos S, DiLillo DJ, Maamary J, Wang TT, Dahan R, et al. Type I and type II Fc receptors regulate innate and adaptive immunity. Nature immunology. 2014;15(8):707–16.

10. van der Poel CE, Spaapen RM, van de Winkel JG, Leusen JH. Functional characteristics of the high affinity IgG receptor, FcgammaRI. Journal of immunology. 2011;186(5):2699–704.

11. Marquis H. Tissue culture cell assays used to analyze Listeria monocytogenes. Curr Protoc Microbiol. 2006;Chapter 9:Unit 9B 4.

12. Vazquez-Boland JA, Kuhn M, Berche P, Chakraborty T, Dominguez-Bernal G, Goebel W, et al. Listeria pathogenesis and molecular virulence determinants. Clin Microbiol Rev. 2001;14(3):584–640.

13. Guillet C, Join-Lambert O, Le Monnier A, Leclercq A, Mechai F, Mamzer-Bruneel MF, et al. Human listeriosis caused by Listeria ivanovii. Emerg Infect Dis. 2010;16(1):136–8.

14. Cummins AJ, Fielding AK, McLauchlin J. Listeria ivanovii infection in a patient with AIDS. J Infect. 1994;28(1):89–91.

15. Snapir YM, Vaisbein E, Nassar F. Low virulence but potentially fatal outcome-Listeria ivanovii. Eur J Intern Med. 2006;17(4):286–7.

16. Gouin E, Mengaud J, Cossart P. The virulence gene cluster of Listeria monocytogenes is also present in Listeria ivanovii, an animal pathogen, and Listeria seeligeri, a nonpathogenic species. Infection and immunity. 1994;62(8):3550–3.

17. Glaser P, Frangeul L, Buchrieser C, Rusniok C, Amend A, Baquero F, et al. Comparative genomics of Listeria species. Science. 2001;294(5543):849–52.

18. Lingnau A, Domann E, Hudel M, Bock M, Nichterlein T, Wehland J, et al. Expression of the Listeria monocytogenes EGD inlA and inlB genes, whose products mediate bacterial entry into tissue culture cell lines, by PrfA-dependent and -independent mechanisms. Infection and immunity. 1995;63(10):3896–903.

19. Milohanic E, Glaser P, Coppee JY, Frangeul L, Vega Y, Vazquez-Boland JA, et al. Transcriptome analysis of Listeria monocytogenes identifies three groups of genes differently regulated by PrfA. Mol Microbiol. 2003;47(6):1613–25.

20. Gaillard JL, Berche P, Frehel C, Gouin E, Cossart P. Entry of L. monocytogenes into cells is mediated by internalin, a repeat protein reminiscent of surface antigens from gram-positive cocci. Cell. 1991;65(7):1127–41.

21. Dramsi S, Biswas I, Maguin E, Braun L, Mastroeni P, Cossart P. Entry of Listeria monocytogenes into hepatocytes requires expression of inIB, a surface protein of the internalin multigene family. Mol Microbiol. 1995;16(2):251 –61.

22. Hamon MA, Ribet D, Stavru F, Cossart P. Listeriolysin O: the Swiss army knife of Listeria. Trends Microbiol. 2012;20(8):360–8.

23. Smith GA, Marquis H, Jones S, Johnston NC, Portnoy DA, Goldfine H. The two distinct phospholipases C of Listeria monocytogenes have overlapping roles in escape from a vacuole and cell-to-cell spread. Infection and immunity. 1995;63(11):4231–7.

24. Poyart C, Abachin E, Razafimanantsoa I, Berche P. The zinc metalloprotease of Listeria monocytogenes is required for maturation of phosphatidylcholine phospholipase C: direct evidence obtained by gene complementation. Infection and immunity. 1993;61(4):1576–80.

25. Rajabian T, Gavicherla B, Heisig M, Muller-Altrock S, Goebel W, Gray-Owen SD, et al. The bacterial virulence factor InlC perturbs apical cell junctions and promotes cell-to-cell spread of Listeria. Nat Cell Biol. 2009;11(10):1212–8.

26. Kocks C, Gouin E, Tabouret M, Berche P, Ohayon H, Cossart P. L. monocytogenes-induced actin assembly requires the actA gene product, a surface protein. Cell. 1992;68(3):521 –31.

27. Chico-Calero I, Suarez M, Gonzalez-Zorn B, Scortti M, Slaghuis J, Goebel W, et al. Hpt, a bacterial homolog of the microsomal glucose-6-phosphate translocase, mediates rapid intracellular proliferation in Listeria. Proceedings of the National Academy of Sciences of the United States of America. 2002;99(1):431–6.

28. Pillich H, Puri M, Chakraborty T. ActA of Listeria monocytogenes and Its Manifold Activities as an Important Listerial Virulence Factor. Curr Top Microbiol Immunol. 2017;399:113-32.

29. Domann E, Wehland J, Rohde M, Pistor S, Hartl M, Goebel W, et al. A novel bacterial virulence gene in Listeria monocytogenes required for host cell microfilament interaction with homology to the proline-rich region of vinculin. The EMBO journal. 1992;11 (5):1981–90.

30. Lauer P, Chow MY, Loessner MJ, Portnoy DA, Calendar R. Construction, characterization, and use of two Listeria monocytogenes site-specific phage integration vectors. Journal of bacteriology. 2002;184(15):4177–86.

31. Gouin E, Dehoux P, Mengaud J, Kocks C, Cossart P. iactA of Listeria ivanovii, although distantly related to Listeria monocytogenes actA, restores actin tail formation in an L. monocytogenes actA mutant. Infection and immunity. 1995;63(7):2729–37.

32. Gerstel B, Grobe L, Pistor S, Chakraborty T, Wehland J. The ActA polypeptides of Listeria ivanovii and Listeria monocytogenes harbor related binding sites for host microfilament proteins. Infection and immunity. 1996;64(6):1929–36.

33. Lecuit M, Ohayon H, Braun L, Mengaud J, Cossart P. Internalin of Listeria monocytogenes with an intact leucine-rich repeat region is sufficient to promote internalization. Infection and immunity. 1997;65(12):5309–19.

34. Travier L, Lecuit M. Listeria monocytogenes ActA: a new function for a 'classic' virulence factor. Curr Opin Microbiol. 2014;17:53-60.

35. Skoble J, Portnoy DA, Welch MD. Three regions within ActA promote Arp2/3 complex-mediated actin nucleation and Listeria monocytogenes motility. The Journal of cell biology. 2000;150(3):527–38.

36. Pistor S, Grobe L, Sechi AS, Domann E, Gerstel B, Machesky LM, et al. Mutations of arginine residues within the 146-KKRRK-150 motif of the ActA protein of Listeria monocytogenes abolish intracellular motility by interfering with the recruitment of the Arp2/3 complex. J Cell Sci. 2000;113 (Pt 18):3277-87.

37. Welch MD, Rosenblatt J, Skoble J, Portnoy DA, Mitchison TJ. Interaction of human Arp2/3 complex and the Listeria monocytogenes ActA protein in actin filament nucleation. Science. 1998;281(5373):105–8.

38. Zalevsky J, Grigorova I, Mullins RD. Activation of the Arp2/3 complex by the Listeria acta protein. Acta binds two actin monomers and three subunits of the Arp2/3 complex. The Journal of biological chemistry. 2001;276(5):3468–75.

39. Lauer P, Theriot JA, Skoble J, Welch MD, Portnoy DA. Systematic mutational analysis of the amino-terminal domain of the Listeria monocytogenes ActA protein reveals novel functions in actin-based motility. Mol Microbiol. 2001;42(5):1163–77.

40. Smith GA, Theriot JA, Portnoy DA. The tandem repeat domain in the Listeria monocytogenes ActA protein controls the rate of actin-based motility, the percentage of moving bacteria, and the localization of vasodilator-stimulated phosphoprotein and profilin. The Journal of cell biology. 1996;135(3):647–60.

41. Van Troys M, Lambrechts A, David V, Demol H, Puype M, Pizarro-Cerda J, et al. The actin propulsive machinery: the proteome of Listeria monocytogenes tails. Biochemical and biophysical research communications. 2008;375(2):194–9.

42. Suarez M, Gonzalez-Zorn B, Vega Y, Chico-Calero I, Vazquez-Boland JA. A role for ActA in epithelial cell invasion by Listeria monocytogenes. Cellular microbiology. 2001;3(12):853–64.

43. Alvarez-Dominguez C, Vazquez-Boland JA, Carrasco-Marin E, Lopez-Mato P, Leyva-Cobian F. Host cell heparan sulfate proteoglycans mediate attachment and entry of Listeria monocytogenes, and the listerial surface protein ActA is involved in heparan sulfate receptor recognition. Infection and immunity. 1997;65(1):78–88.

44. Poussin MA, Goldfine H. Evidence for the involvement of ActA in maturation of the Listeria monocytogenes phagosome. Cell Res. 2010;20(1):109–12.

45. Yoshikawa Y, Ogawa M, Hain T, Yoshida M, Fukumatsu M, Kim M, et al. Listeria monocytogenes ActA-mediated escape from autophagic recognition. Nat Cell Biol. 2009;11(10):1233– 40.

46. Mitchell G, Ge L, Huang Q, Chen C, Kianian S, Roberts MF, et al. Avoidance of autophagy mediated by PlcA or ActA is required for Listeria monocytogenes growth in macrophages. Infection and immunity. 2015;83(5):2175–84.

47. Travier L, Guadagnini S, Gouin E, Dufour A, Chenal-Francisque V, Cossart P, et al. ActA promotes Listeria monocytogenes aggregation, intestinal colonization and carriage. PLoS Pathog. 2013;9(1):e1003131.

48. Welch MD, Way M. Arp2/3-mediated actin-based motility: a tail of pathogen abuse. Cell Host Microbe. 2013;14(3):242–55.

49. Kreft J, Dumbsky M, Theiss S. The actin-polymerization protein from Listeria ivanovii is a large repeat protein which shows only limited amino acid sequence homology to ActA from Listeria monocytogenes. FEMS Microbiol Lett. 1995;126(2):113–21.

50. Chakraborty T, Hain T, Domann E. Genome organization and the evolution of the virulence gene locus in Listeria species. Int J Med Microbiol. 2000;290(2):167–74.

51. Moors MA, Levitt B, Youngman P, Portnoy DA. Expression of listeriolysin O and ActA by intracellular and extracellular Listeria monocytogenes. Infection and immunity. 1999;67(1):131–9.

52. Freitag NE, Jacobs KE. Examination of Listeria monocytogenes intracellular gene expression by using the green fluorescent protein of Aequorea victoria. Infection and immunity. 1999;67(4):1844–52.

53. Shetron-Rama LM, Marquis H, Bouwer HG, Freitag NE. Intracellular induction of Listeria monocytogenes actA expression. Infection and immunity. 2002;70(3):1087–96.

54. Renzoni A, Cossart P, Dramsi S. PrfA, the transcriptional activator of virulence genes, is upregulated during interaction of Listeria monocytogenes with mammalian cells and in eukaryotic cell extracts. Mol Microbiol. 1999;34(3):552–61.

55. Schoggins JW, Wilson SJ, Panis M, Murphy MY, Jones CT, Bieniasz P, et al. A diverse range of gene products are effectors of the type I interferon antiviral response. Nature. 2011;472(7344):481–5.

56. Simon R, Priefer U, Pühler A. A Broad Host Range Mobilization System for In Vivo Genetic Engineering: Transposon Mutagenesis in Gram Negative Bacteria. Nature Biotechnology. 1983;1(9):784–91.

57. Cheng LW, Portnoy DA. Drosophila S2 cells: an alternative infection model for Listeria monocytogenes. Cellular microbiology. 2003;5(12):875–85.

